# Microenvironment-contextual cell signaling is attenuated with age

**DOI:** 10.1101/684977

**Authors:** Henriette C. Ertsås, Mark A. LaBarge, James B Lorens

## Abstract

Post-menopausal women are more prone to breast cancer than younger women. The increased frequency of age-related breast cancers is likely due to interactions between acquired mutations and age-dependent epigenetic changes that affect mammary epithelial lineage fidelity. We hypothesized that the aging process fundamentally affects how human mammary epithelial cells (HMEC) respond to microenvironmental signals, resulting in increased susceptibility to oncogenic transformation. In order to measure microenvironmental cell signaling in normal finite lifespan HMEC, we applied a novel microsphere-based flow cytometry technology. The microsphere cytometry allows multiparametric single cell quantification of signaling pathway activity and lineage-specific marker expression in cells adhered to surface-functionalized microspheres that mimic specific microenvironments. Using this approach, we analyzed age-dependent changes in human mammary myoepithelial and luminal epithelial cells exposed to different ECM and growth factors. We found that ECM–mediated MAP kinase and PI3 kinase activation levels in HMEC were attenuated with age. Older luminal cells displayed higher surface integrin levels consistent with acquired basal identity, albeit with decreased integrin activation and increased Src-signaling relative to myoepithelial cells. We show that the diminished signaling magnitude in HMEC from older women correlated with reduced probability of activating oncogene-induced senescence. We propose that age-related changes in ECM-mediated epithelial cell regulation may impair protective tumor suppression mechanisms and increase breast cancer susceptibility.

## Introduction

Age is indisputably the highest risk factor for developing cancer. A recent study reported that over half of current adults under the age of 65 years will be diagnosed with cancer at some point in their lifetime, due to increased overall lifespan ^1^. More than 75% of breast cancers occur among woman over the age of 50 ^2^. Accumulation of gene mutations is the predominant model to explain these observations. However, age related breast cancer is not associated with specific mutations and the trend attenuates after the age of 70 ^3^. The mechanisms of aging comprise common intrinsic factors; such as telomere shortening and senescence, and tissue-level factors; such as replacement of epithelial cells with adipocytes and accumulating loci of fibrotic rigidity due to glucose oxidation and subsequent crosslinking of collagen I ^4–7^. The conflation of intrinsic and microenvironmental factors results in significant age-related transcriptional changes ^8,9^. Aging is associated with emergent mammary epithelial cell phenotypes in particular in the luminal compartment. Subsets of luminal cells acquire basal traits that are related to phenotypic changes associated with BRCA1 mutations ^10,11^. We reasoned that age-emergent HMEC phenotypes will respond differently to microenvironmental cues and affect tumor suppressive senescence barriers. To address this possibility, we investigated normal, pre-stasis, finite lifespan human mammary epithelial cells (HMEC) from reduction mammoplasty, deriving from two groups of women; aged <30(y)ears and >58y. We applied a novel microsphere cytometry method to measure HMEC signaling at the single cell level within heterogeneous cell populations in the context of different extracellular matrix proteins (ECM) and growth factors. We found altered ECM-mediated MAPK and PI3K-AKT pathway signaling with age, and evidence that decreased signaling in aged HMEC reduces the probability of activating tumor suppressive senescence barriers^12^.

## Results

### ECM-mediated signaling was altered in older human mammary epithelial cells

In order to measure ECM-mediated cell signaling responses in adherent HMEC, we applied a novel microsphere cytometry technique ^13^. Microspheres were coated with E cadherin to mimic cell-cell contact ^14^ and extracellular matrix proteins (ECM) (collagen 1 (COL1), fibronectin (FN) or laminin (LAM)) to mimic the extracellular matrix (Fig. 1A), and mixed with low passage (4p) pre-stasis HMEC strains ^15^ derived from reduction mammoplasty or mastectomy tissues from women between the age of 19-91y. HMEC were allowed to adhere to coated microspheres for 2.5 hours prior to treatment and analysis by flow cytometry. DNA staining and side scatter analysis allowed unambiguous identification of single cells bound to individual microspheres (Fig.1B). Human mammary luminal (LEP) and myoepithelial (MEP) cells were distinguished by expression of cell surface markers (MEP:CD10 ^+^CD227^-^; LEP:CD227^+^ CD10^-^). Staining with anti-pERK and anti-pAKT was used to measure MAPK and PI3-AKT pathway activation respectively. Basal phosphorylation levels in cells were established by pretreatment with pERK and pAKT inhibitors (PD98059, wortmannin). Cells with phosphoprotein-levels above the basal phosphorylation levels were defined as responding (Fig.1C). The geometric mean fluorescence intensity (GMFI) value of anti-pERK and anti-pAKT levels in the responding population was used to determine the strength of cell response at different ages (Fig.1C). Linear regression analysis of GMFI of the responding populations as a function of age revealed a reduction in the responses of LEP and MEP to different ECM proteins with age (Fig.1D; Suppl. Fig.1). pERK activation levels were significantly correlated with age in both LEP and MEP when adhered to FN. A similar trend was observed for COL1 and LAM, suggesting that ECM-mediated cell signaling was diminished with age. Notably, the relative percentage of HMEC within the response gate did not change, showing that the attenuated phosphoprotein levels were due to decreased signaling magnitude and not fewer responding cells (Suppl. Fig.2C)

**Fig.1.**
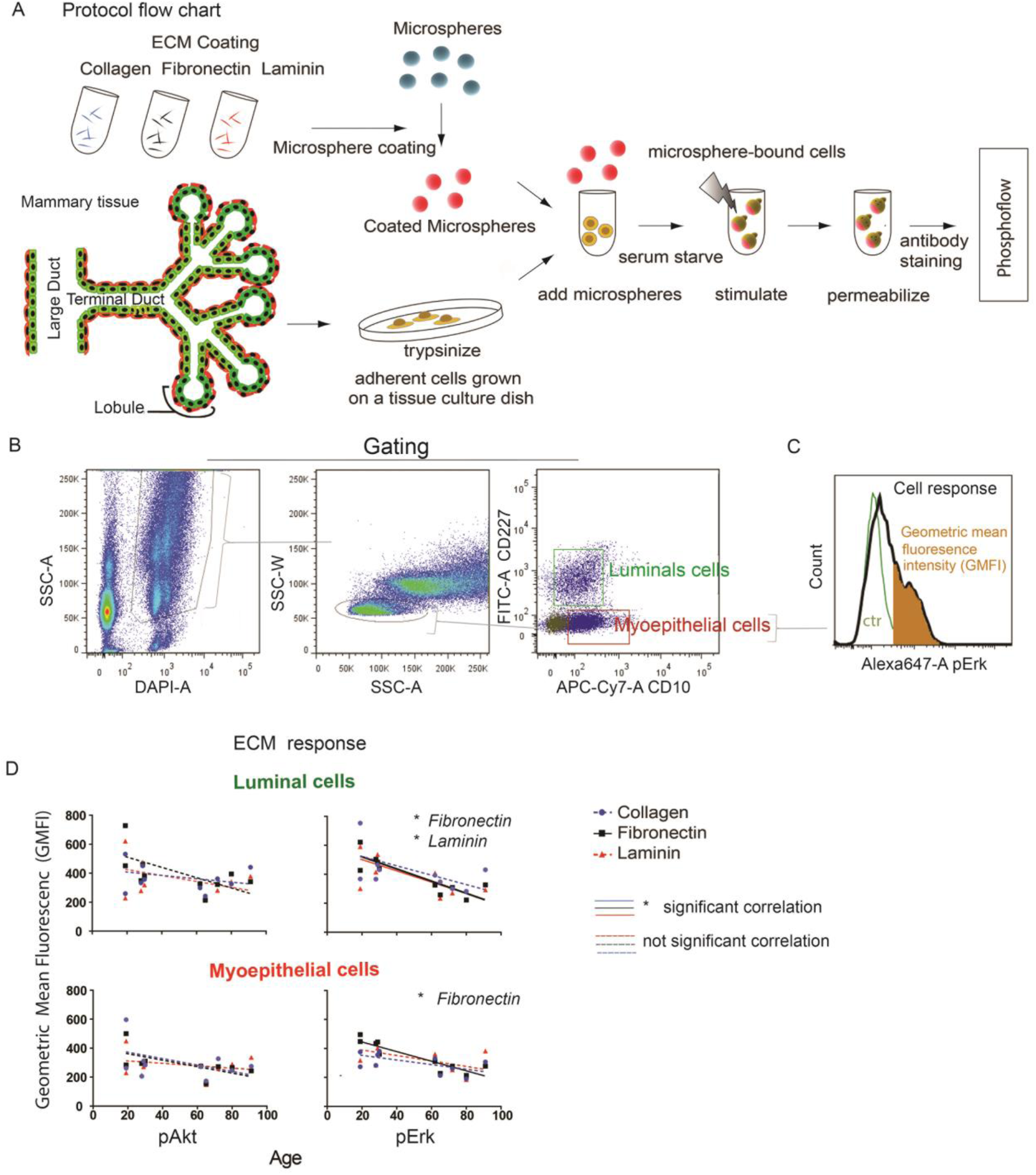
Mammary epithelial cell signaling responses to ECM is attenuated with age. Schematic representation of the microsphere cytometry approach. Primary human mammary epithelial cells (HMEC) obtained from reduction mammoplasty and mastectomy are adhered to ECM-coated microspheres. (B) Flow cytometry gating strategy to resolve single cells adhered to individual microspheres. The two mammary epithelial lineages derived from different compartments of the breast epithelium were analyzed separately: luminal [LEP] (CD10^-^CD227^+^) and myoepithelial [MEP] (CD10^+^CD227^-^). (C) Geometric mean fluorescence intensity measurement of pERK levels in gated microsphere-bound cells. The activation gate (orange) was determined by phosphorylation levels in the presence of a specific kinase inhibitor used to define the negative control (ctr, green). (D) Phospho-protein (pAKT and pERK) analysis of 10 women between the age of 19 and 91 y by HMEC adhered to COL1, FN or LAM-coated microspheres. Linear regression analysis of data shows reduced ECM induced signaling in LEP and MEP as a function of age. GMFI = geometric mean fluorescent intensity. Significant correlations (* P<0.05) are demarcated by solid lines; non-significant correlations are represented by dotted lines.

In order to determine growth factor-mediated AKT and ERK activation in different ECM contexts, we stimulated microsphere-bound HMEC for 20 minutes with complete M87 medium, comprising several growth factors (see experimental methods), and measured the relative change in pAKT and pERK levels for HMEC (Suppl. Fig.2A). AKT and ERK activation in response to the growth factors did not show significant changes with age (Suppl. Fig. 2B). We propose that HMEC exhibit age-dependent responses to different ECM, but that growth factor-mediated activation is age independent.

### Cell responsiveness to ECM is altered with increasing age

In order to further investigate the nature of the ECM response with age, longitudinal cell signaling responses were analyzed in HMEC adhered to ECM-coated microspheres while incubated in M87 complete medium. pERK levels were monitored in HMEC between 0-2 hours following adhesion to ECM-coated microspheres. HMEC displayed a sigmoidal pERK response with the major inflection occurring between 0.5-1 hr following ECM adhesion (Fig.2). Nonlinear regression analyses of normalized data with variable slope revealed a significant delay in pERK activation in HMEC from older women following binding to COL1 or LAM (Fig.2A). Cells bound to FN showed no differences in pERK activation kinetics (Fig.2B). In contrast, pAKT induction in adhered HMEC remained relatively stable at the measured timepoints (Suppl. Fig.3A)

**Fig.2.**
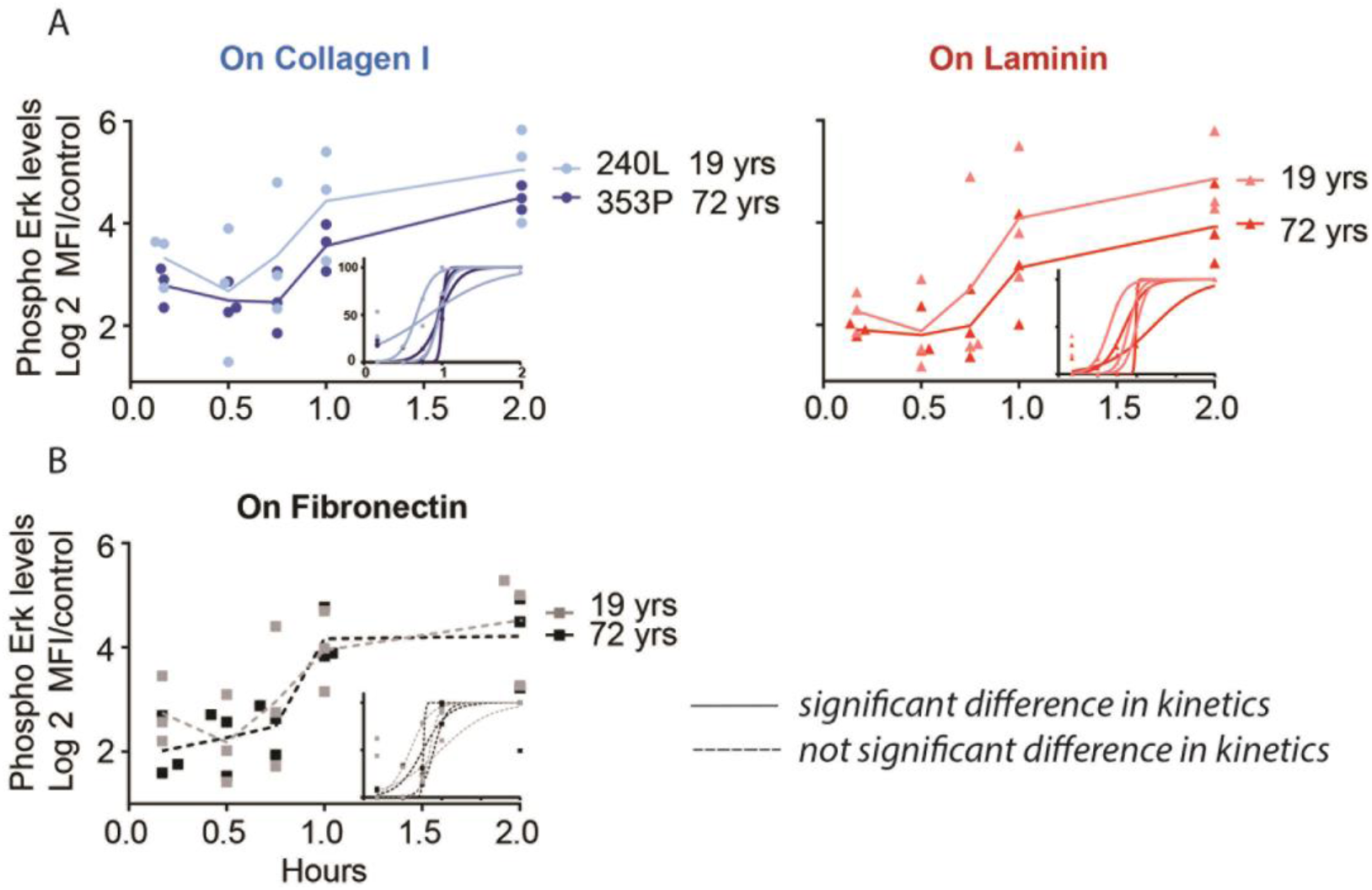
Contextual cell responsiveness is altered with increasing age. Kinetic analysis of pERK was performed on HMEC derived from women 19 and 72 y while bound to ECM-coated microspheres (COL1, FN and LAM) for a 2-hour period (n=3). Control samples stained with secondary antibody only, represent background. Plotted numbers are log2 ratios of sample over background. Nonlinear regression variable-slope analysis is shown in the lower right corner of each plot, a data point represents the percentage value of the maximum and minimum phospho-level within the 2-hour period. HMEC-data from young women are indicated by a lighter color, while a darker color represents HMEC from older women. (A) A significant delay in pERK responsiveness was found when comparing HMEC adhered to COL1 and LAM (P = 0.03). P-value was calculated by student’s one tailed t-test comparing LogEC50 values. (B) HMEC bound to FN did not show significant difference in pERK responsiveness with age (significance difference = solid line, not significant = dotted line).

Given that MEP reside in the basal compartment, we monitored pERK and pAKT levels in the MEP subpopulation between 0-9 hr post-ECM adhesion (Suppl. Fig.3B). The MEP from younger women reached maximum pERK levels between 2-4 hours on different ECM. MEP from older women displayed comparatively delayed and lower activation levels (Suppl. Fig.3Bi)). Interestingly, MEP from the older women appeared to reach maximal pAKT levels on COL1 and FN earlier than the young (Suppl. Fig.3Bii). Overall, HMEC from older women displayed altered ECM responsiveness.

### Integrin activation by ECM decreases with age

Next, we asked if HMEC cell adhesion to different ECM was affected by age. Interestingly, overall cell adhesion increased with age, both in percentage of adhered cells within a given time and as the rate of adhesion (Suppl. Fig.4Aii and B). This is consistent with previous results showing that HMEC aging is associated with increased basal characteristics^16^. Congruently, total surface integrin levels on HMEC increased with age (Suppl. Fig.5). We further determined whether there were changes in the relative proportion of activated integrins (ITG) in HMEC adhered to ECM-coated microspheres by measuring ITGβ1 and ITGβ4 phosphorylation and downstream pSrc levels (Fig.3). The ratio of integrin activation in LEP to MEP was used to determine the extent of basal identity as a function of age, where values >0 were defined as more basal-like, and <0 more luminal-like in a COL1 context. This analysis indicated that LEP acquired a significantly greater basal identity with increasing age (Fig.3Bii). This was significant for Src activation in a context of COL1 as a function of age, with a similar trend found for all integrins measured. This relative increase in integrin activation levels was mainly due to reduced integrin phosphorylation in the MEP subpopulation. However, taking into account the increase in overall integrin expression (Suppl. Fig.5B), we found that relative integrin activation decreased with age in both LEP and MEP (Fig.3C), with a significant decrease for ITGβ4 phosphorylated on Y1510. Taken together our findings suggested that LEP acquire a basal identity with age, while relative integrin activation in a context of COL1 decreased in both LEP and MEP with age.

**Fig.3.**
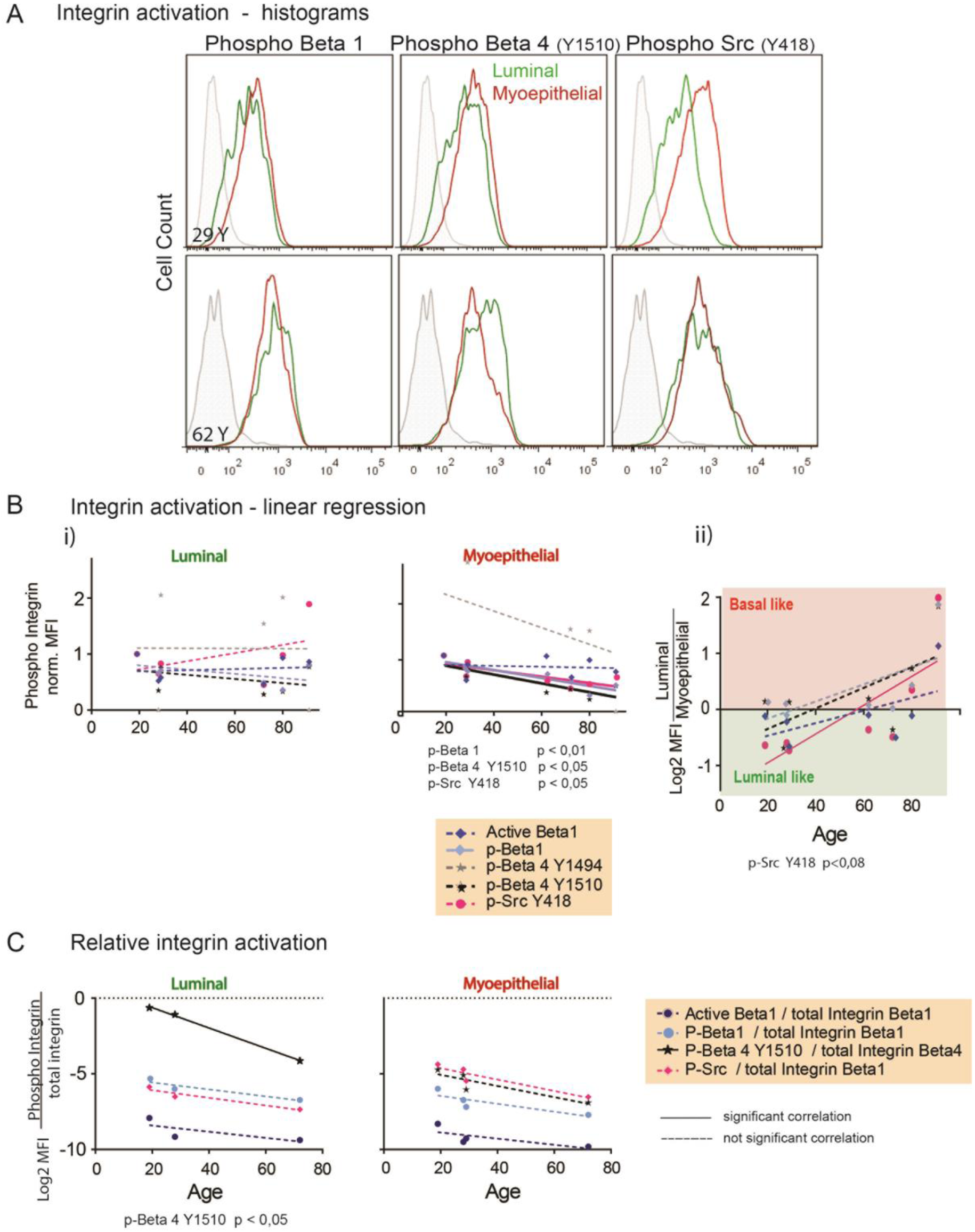
Integrin activation distinguishes luminal and myoepithelial cells during aging. HMEC from 8 women between 19 and 91 y were adhered to COL1-coated microspheres and integrin activation in the LEP and MEP subpopulations was measured with by Phosflow cytometry of Src (pY418), ITGβ1 (pT789) and ITGβ4 (pY1494, pY1510). An antibody recognizing the active confirmation of ITGβ1 was also included. (A) Representative histograms of microsphere cytometry analysis of LEP and MEP subpopulations from HMEC derived from women 29 and 62 y, adhered to FN-coated microspheres. (B) Integrin activation analyzed by linear regression as a function of age. i) Data normalized to the youngest HMEC, each data point represents MFI (median fluorescence intensity) of sample / MFI of HMEC (19y). ii) Log2 ratio of LEP vs MEP. LEP trends towards a more basal identity. Values >0 were defined as basal-like. (C) HMEC from 3 women between 19 and 72 y were adhered to COL1-coated microspheres, and the resulting integrin activation was measured by microsphere cytometry. Relative activation presented as the log2 ratio of phosphorylated integrin over total integrin. (significance difference = solid line, not significant = dotted line).

### Age-related susceptibility of HMEC to oncogene induced senescence

We questioned whether the observed age-related changes in HMEC responsiveness to ECM could affect tumor suppressive mechanisms and alter susceptibility to oncogenic transformation. Oncogene induced senescence (OIS) is a key protective mechanism triggered by abnormal cell signaling^17^. We investigated by the use of a constitutive active EGFR mutant oncogene (EGFR^Del19^), whether there were age-related differences in OIS induction in pre-stasis HMEC. EGFR^Del19^ or empty retroviral vectors was introduced into pre-stasis HMEC. EGFR^Del19^ expressing cells were sorted 48 hours later to ensure uniform EGFR expression (Suppl. Fig.6A), and EGFR expression was followed during cell passaging (Suppl. Fig.6B). Cells with empty retroviral vector were selected by antibiotics. EGFR^Del19^ oncogene dependency in transduced cells was confirmed by erlotinib sensitivity (Suppl. Fig.7). Cells were allowed to adhere to ECM-coated microspheres, followed by cell signal analysis by microsphere cytometry (Fig.4A). Serum-deprivation was applied to secure mutant EGFR signaling (open dots); some sample pairs had additional EGF treatment (closed dots). Isogenic immortalized and tumorigenic cell lines were included for comparison. Data was plotted as the log2 value of signaling in EGFR^Del19^-overexpressing cells versus empty retroviral vector-transduced cells as a function of age. Linear regression analysis revealed a reduction of EGFR^Del19^-induced pERK and pAKT with HMEC age. We quantified senescence by measuring β-galactosidase activity with X-Gal (blue) in EGFR^Del19^-expressing HMEC from a 28 and 58y woman (Fig.4B). The HMEC from younger women showed large vacuoles and reduced proliferation rapidly after expression of EGFR^Del19^. In contrast, the HMEC from the older women displayed less β-galactosidase activity, and demonstrated a mesenchymal morphology. These cells proliferated for several passages beyond the younger EGFR^Del19^-transduced HMEC (Fig.4C). Collectively, these results suggest that triggering of OIS by oncogenic EGFR expression was reduced in HMEC with age, consistent with an increased susceptibility to oncogenic transformation.

**Fig.4.**
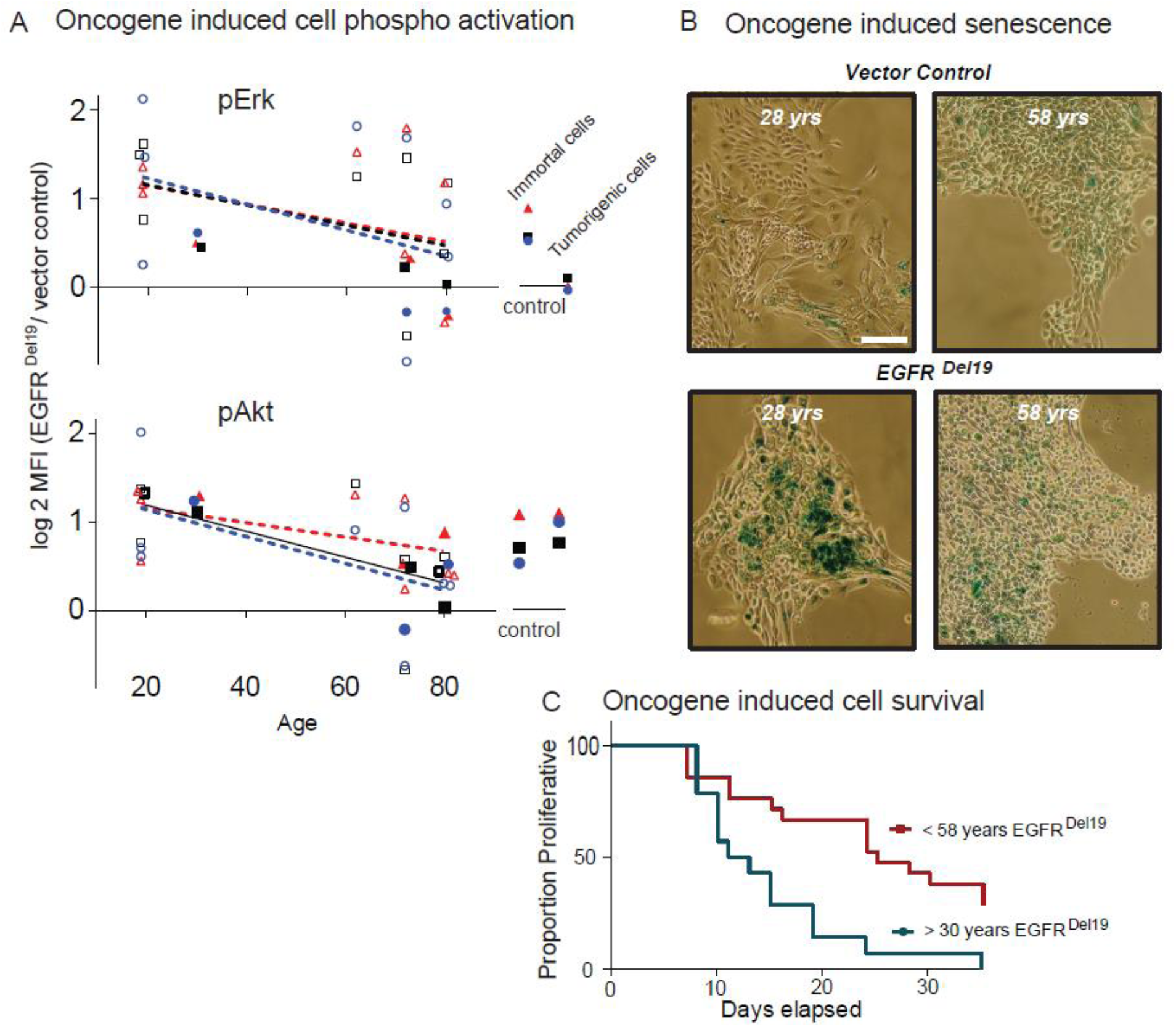
Older HMEC show reduced oncogene-induced senescence (OIS). (A) Linear regression analysis of EGFR^Del19^ – induced ERK and AKT activation as a function of age. Data points represent the log2 ratio of phospho-levels of paired samples of EGFR^Del1-^ expressing and control HMEC from 6 different women age 19-80, with (filled dots) or without (open dots) additional EGF treatment. The experiment was performed in three different ECM contexts, COL1, FN and LAM. Cells from the HMEC progression series of increasing tumorigenicity were added as controls (immortal and tumorigenic). (significance difference (P < 0,05) = solid line; not significant = dotted line). (B) Phase-contrast images of the levels of cell senescence in women (28 and 58 y) following EGFR^Del19^ overexpression. Senescence is detected by the presence of β-galactosidase activity. Bar: 100 µm. (C) Kaplan-Meyer analysis of survival time of EGFR ^Del19^ transduced cells from 7 young and 8 older women. Period of survival of cells is stated in days.

## Discussion

Aging is associated with both intrinsic and extrinsic changes in the mammary epithelium. The post-menopausal breast microenvironment is characterized by basal membrane degradation, an increasingly collagenous stroma and gradual replacement of a senescent epithelia with soft adipose and inflammatory cells ^18^. Aging HMEC, originally sequestered by a basal membrane consisting of LAM and COL4, become exposed to different ECM components such as FN and COL1, normally only encountered during development, wound healing and inflammation ^19,20^.

We show that PI3K and MAPK signaling following HMEC cell adhesion to LAM and FN is attenuated with age (Fig.1, Fig.2). This reduced responsiveness correlates with changes in integrin expression and function (Suppl. Fig.5, Fig.3). Our single cell-level microsphere cytometry analysis revealed that this was due to reduced signal magnitude and not a reduction in the number of responding cells (Suppl. Fig.2C). HMEC from younger women demonstrated distinctive pERK responses to COL1, FN or LAM, while responses in HMEC from older women were largely independent of ECM type (Suppl. Fig.1). We also detected lower levels of ITGβ4 activation in LEP from older women (Fig.3C), indicating a change in ECM-mediated cell signaling with age.

LEP normally have lower integrin expression than MEP which reside in the basal compartment. ITGα2 and ITGβ1 were the most prevalent on both LEP and MEP (Suppl. Fig.5B). However, we observed increased expression of ITGα2, ITGβ1, ITGα6 and ITGβ4 integrins on LEP with increasing age. This is consistent with earlier observations that LEP acquire basal traits with age^16^. It may be related to a concomitant reduction in the total number of MEP and structural alternations in the basal membrane with age that disrupt the epithelial bilayer and give LEP abnormal access to the ECM^21^. This breakdown in normal tissue architecture is thus associated with age-emergent cellular phenotypes with altered cell signaling responses. Oncogenic mutations in healthy cells that enhance cell signaling normally induce senescence (OIS)^22^. We show that the attenuated cell signaling responses in HMEC derived from older women can subvert this important tumor defense mechanism. ERK and AKT activation levels generated by an oncogenic form of EGFR were reduced in older HMEC (Fig.4). This surprising result indicates that HMEC, with age-induced epigenetic alterations, may be less prone to OIS and thus more likely to tolerate oncogenic mutations. This raises the interesting possibility that drugs targeting epigenetic regulators could be used in breast cancer prevention ^23–25^.

## Materials and Methods

### Coating of Microspheres with Extracellular Matrix Proteins

VARIAN PL-Microspheres SuperCarbonyl White poly(styrene-co-methancrylic acid) 20 μm in diameter (Batch CD185, Agilent Technologies) were coated as described ^13^. E-Cadherin (1 µg mL^-1^, Sino-Biological Inc. #10204-H08H) was added to the coating suspension..

### Cell culture

Finite life span mammary epithelial cells were obtained from women having undergone reduction mammoplasty, and generously provided from the HMEC biobank^26^. Cells were cultured in M87A complete medium as described ^13^. Cells were not cultured beyond passage 4. Organoids were cryostored, then thawed and analyzed directly without culturing. Phoenix retroviral packaging cells were grown in DMEM supplemented with 1mM L-glutamine, 10% fetal bovine serum, 100U mL^-1^ Penicillin and 100µg mL^-1^ streptomycin^27^.

Immortalized and oncogene-expressing cells were grown in M87 complete medium supplemented with antibiotics in order to maintain selection during culturing. Hygromycin (5 µg mL^-1^, Sigma-Aldrich #H3274) enriched for cMyc expressing cells, blasticidin (5 µg mL^-1^, Sigma-Aldrich #15205) maintained Neu/Her2 expression, and pyromycin (0.5 µg mL^-1^, Sigma-Aldrich #P8833) maintained EGFR overexpression in EGFR^Del19^ transduced cells.

### Microsphere cytometry protocol

Cultured cells were washed twice with PBS and treated with low concentration trypsin (0.05% trypsin, 0.02% EDTA in PBS). Microsphere adhesion was performed as described ^13^. Cell - microsphere incubation time varied between 2-6 hours depending on cell type and choice of ECM coating. Microsphere-bound cells were serum starved for 35 min. prior to stimulation with M87A complete medium for 20 min. (Fig.1) or EGF 20 ng mL^-1^ for 10 min. (Fig.4A). However, EGFR-transduced microsphere-bound cells were mainly cultured in serum-free medium throughout the experiment (Fig.4A). Kinetic analysis of microsphere-bound cell activation, and measurement of integrin expression and activated integrin levels were performed in M87 complete medium (Fig.2 and Fig.3). Pre-treatment (2hr) with MAPK inhibitor PD98059 (20 μM, Cell Signaling #9900) and PI3K inhibitor wortmannin (0.5 μM, Sigma-Aldrich #W1628) was used to set a gate defining the positive population of cells with active ERK or active AKT respectively (Fig.1 and 4). Cell-microsphere complexes were fixed and permeabilized as described^13^.

### Flow cytometry analysis

Fixed, permeabilized cell-microsphere complexes were stained as described ^28^. We used the following rabbit primary antibodies diluted in 1% bovine serum albumin in PBS as a blocking buffer: anti-pERK, (1/200, Cell Signaling #4370), anti-pAKT (1/200, Cell Signaling #4060), anti-pY1494 ITGβ4 (1/100, AbCam #29043), anti-pY1510 ITGβ4 (1/100, AbCam # 63546), anti-pY418 Src (1/100, Invitrogen #44660G), anti-pT789 ITGβ1 (1/50, AbCam #138432); FITC conjugated antibody against active [12G10] ITGβ1 (1/50, AbCam #15002).

The following conjugated murine antibodies against integrins were diluted 1/100 and were purchased from Biolegend: PE anti-ITGα2 (#359307), PE anti-ITGα5 (#328009), PE anti-ITGαV (#327909), PE anti-ITGα6 (#313611) PE anti-ITGβ1 (#303003), PE anti-ITGβ4 (#327807). PE anti-EGFR from Biolegend was also diluted 1/100 (#352904). The following lineage specific antibodies were used to separate LEP, MEP and progenitor subpopulations: FITC anti-CD227 (MUC-1) (1/200, #559774) from BD biosciences and APC-Cy7 anti-CD10 (1/100, #312212) PE anti-CD117 (1/100, #313210) from Biolegend.

The secondary antibody goat anti-rabbit Alexa Fluor 647 (#A21244) from Invitrogen was used at a dilution of 1/2500 (0.8 µg/mL). DAPI (Sigma-Aldrich #D9542) or Hoechst (Sigma-Aldrich #B2261) was introduced together with the secondary antibody in a concentration of 1 µg mL^-1^ and 0.6 µg mL^-1^, respectively. All centrifugation steps involving microspheres were spun at 390g for 2 min. Cell suspensions without microspheres were spun at 200g for 5 min. Duration of staining was 30 min. in room temperature for primary antibodies, and at 4°C for cell surface antibodies and secondary antibodies. Samples were analyzed on BD FACS Vantage or BD LSRFortessa. Cell sorting of EGFR^Del19^-transduced cells or CyclinD1-transduced cells was done with BD FACS ARIA and FACS Vantage respectively. All Cytometers were purchased from BD Biosciences. Data was analyzed with Flow Jo (Tree Star Inc.) for the making of histograms.

Staining of microsphere bound cells was performed in two steps: 1) staining of intracellular epitopes with primary antibodies against phosphorylated proteins (pERK, pAKT, pITGβ1, pSrc, pITGβ4); 2) Staining of cell surface markers (CD227, CD10, CD117) combined with secondary antibodies against the rabbit primary antibodies used in the first step, in addition to a DNA stain.

### Kinetic signal transduction analysis

The kinetic experiments were initiated in M87 complete medium. Microsphere-bound cells were treated according to protocol described above. Cell-microsphere complexes were collected at 10, 30, 45, 60, 120, 240 and 540 min. by adding PFA and methanol as described above. Median fluorescence intensity was plotted according to increasing period of ECM/microsphere exposure. Kinetic signaling data was normalized by calculating log2 of the ratio of sample fluorescence against background fluorescence. Background was defined as the fluorescence of a control sample stained with secondary antibody only. In order to investigate significant differences in kinetics, we recalculated the sample values into ratio values where minimum value of the experiment data was set to 0 and maximum value was set to 100. From this set-up EC50 values were obtained and significant differences in kinetics could be detected. Adhesion kinetics was plotted as Hill’s curve, and area under the curve was calculated to investigate significant differences in cell-microsphere adhesion. Experiments were performed in three parallels. Samples were run on BD LSRFortessa (BD Biosciences).

### Extraction of cells from organoids

Cryostored organoids were thawed in a 37°C water bath and transferred to a 50 mL Falcon tube, followed by centrifugation at 450g for 5 min. in a volume of 30 mL PBS. Supernatant was aspirated and the pellet gently resuspended in 3 mL 0.25% Trypsin/EDTA. The suspension was placed on an orbital shaker for 10 min. at room temperature, followed by vigorous shaking for 30 seconds. 10 mL of medium was added to neutralize trypsin and the suspension was filtered through multiple 70 µm strainers until no visible debris was found in the filtrate. Filtrate was pelleted by 600g centrifugation for 5 min. and stained directly for integrin analysis.

### Erlotinib-treatment

The following concentrations of Erlotinib were tested: 0, 1, 3, 10, 30, 100, 300, 1000, 3000 and 10 000 nM. 2000 cells were seeded into each well in a 96 well plate with 8 parallels for each concentration of inhibitor. Survival after 3 days of treatment was measured by the use of Resazurin (Promega #G8081). Cells were incubated with Resazurin for 3 hours. Pink Resorufin is the result of cell metabolism (emission maximum at 580nm), and can be detected in a fluorometer (Perkin Elmer, Wallac 1420 Mulitlabel Counter). 100% cell survival was defined as the level of Resorufin in samples treated with 0nM Erlotinib. Experiment was performed as described in CellTiter-Blue®Cell Viability Assay protocol from Promega.

### Senescence associated β-Galactosidase activity

β-galactosidase assay was performed as described ^29^, with a buffer pH at 6 to selectively stain only senescent cells growing in cell culture dishes. Cells were washed 3X with PBS before adding fixative. Microsphere-bound cells were stained overnight, washed and images were retrieved in a phase-contrast microscope.

### Statistics

All statistical tests were performed in GraphPad Prism 6

## Acknowledgements

We thank Mark La Barge for welcoming us into his lab, and for providing insight and material for the research on the topic of aging. We thank Fanny Pellisier for helpful discussion around additional aspects of this project; Dr. Martha Stampfer for contributing breast tissue from collected tissue samples, and instructions in how to care for these cells; Michelle Scott for invaluable technical assistance in flow cytometry and James Garbe for sharing with us methods to make the cell thrive. H.E was supported by a Norwegian Cancer Society predoctoral fellowship.

---

**Suppl. Fig.1.**
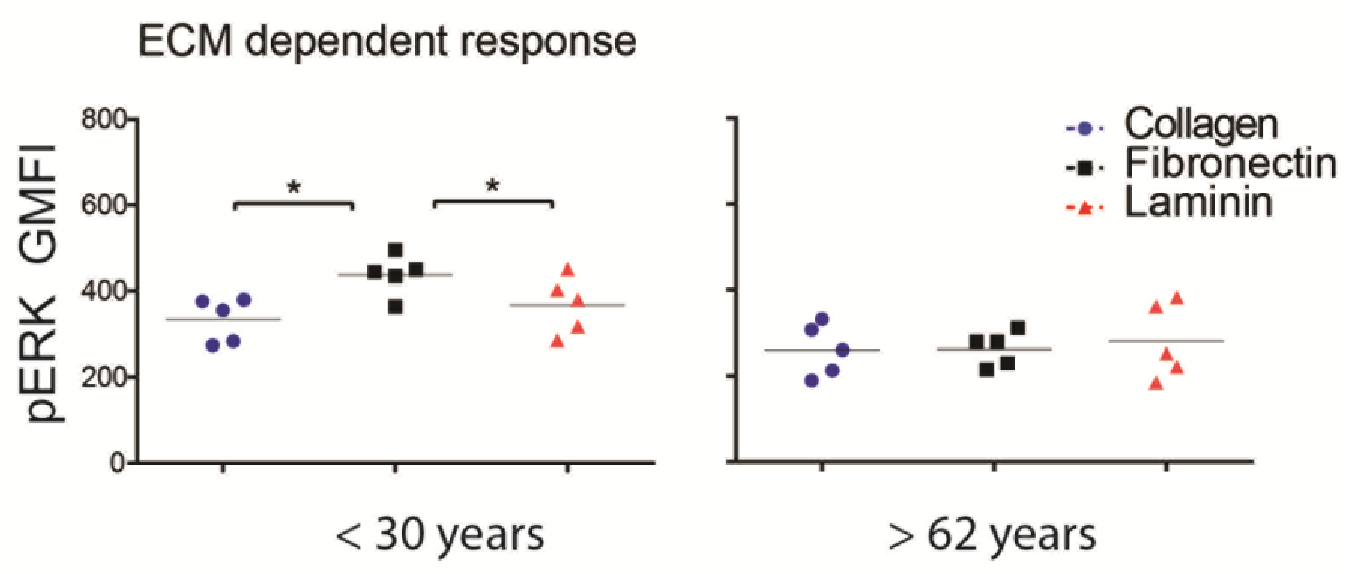
Mammary epithelial cell pERK responsiveness to fibronectin is reduced with age. pERK levels in HMEC from women < 30y and > 62y adhered to COL1, FN or LAM were measured. HMEC from 10 women between the age of 19 and 91 were included. The pERK GMFI (geometric mean fluorescence intensity) values compared were defined according to description in Fig.1 (C). HMEC from younger women showed a stronger pERK responsiveness when bound to FN than to COL1 or LAM. *P<0.5

**Suppl. Fig.2.**
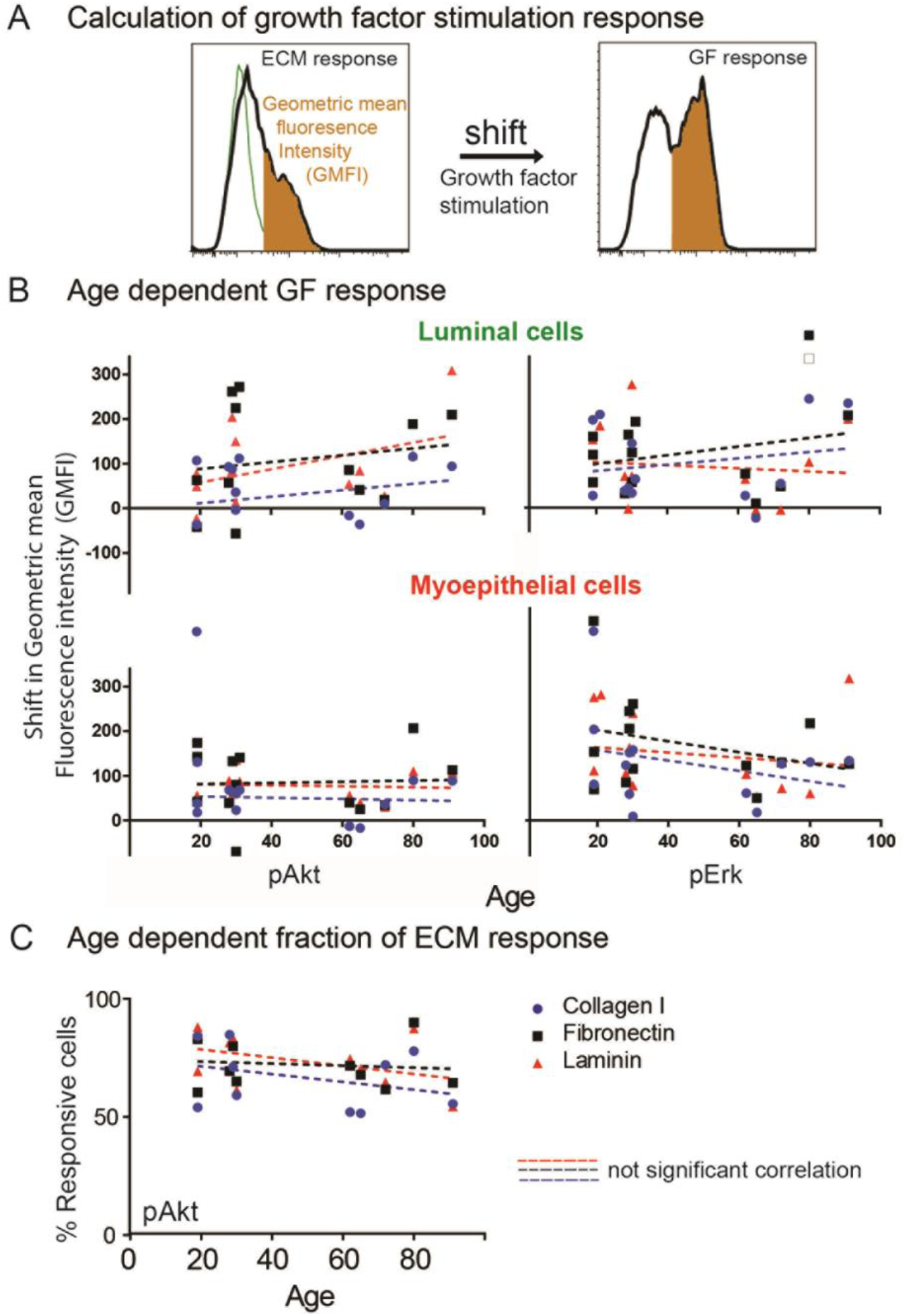
Mammary epithelial cell responsiveness to growth factor stimulation is independent of age. (A) Growth factor (GF)-induced responses are defined as the shift in GMFI of the responding population (in orange) upon GF stimulation. The GMFI levels measured in microsphere-bound cells in the absence of growth factors was subtracted to calculate GF induced shift. (B) Phospho-protein (pAKT and pERK) analysis of low passage human mammary epithelial cells (HMEC) from 10 women between the age of 19 and 91 y adhered to COL1, FN or LAM-coated microspheres. Linear regression analysis of shift in GFMI upon growth factor stimulation in LEP and MEP as a function of age. (C) The percentage of cells defined as responsive to ECM did not change as a function of age. Non-significant correlations are represented by dotted lines. *P<0.5

**Suppl. Fig.3.**
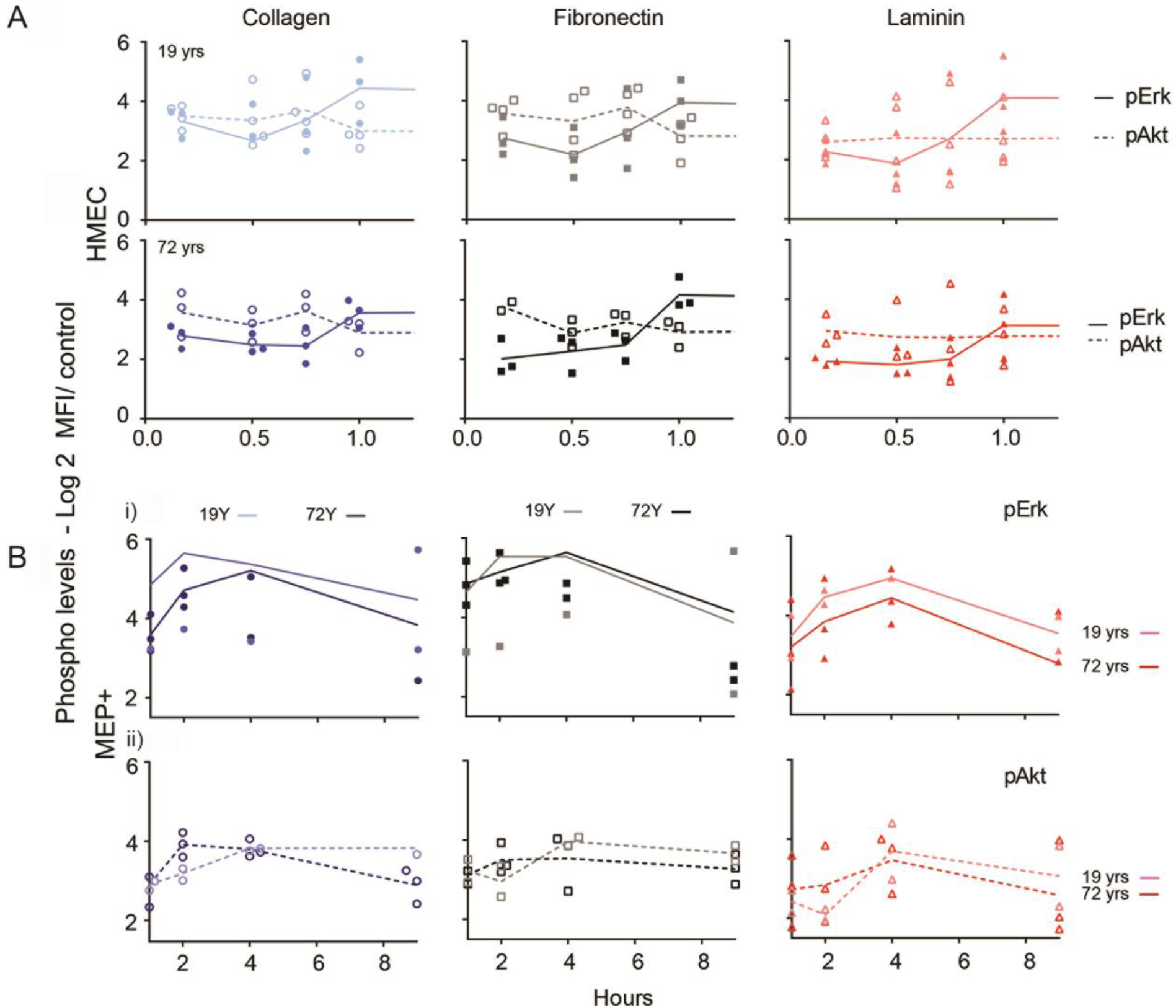
Crosstalk between the pathways of PI3 kinase and MAP kinase is attenuated with age. Analysis of ECM-mediated signaling in HMEC from women 19 and 72 y (n=3 for each woman) during a 9-hour period following adhesion to ECM-coated microspheres (COL1, FN and LAM). Solid lines represent pERK, dotted lines represent pAKT. Data of HMEC from a young woman are indicated by a lighter color, while a darker color represents HMEC from an older woman. (A) pERK and pAKT levels monitored between 0-1,25 hr. (B) MEP+ gated cells monitored between 0-9 hr. i) pERK and ii) pAKT control samples were stained with secondary antibody only and represent background levels. Plotted numbers are log2 ratios of sample over background staining.

**Suppl. Fig.4.**
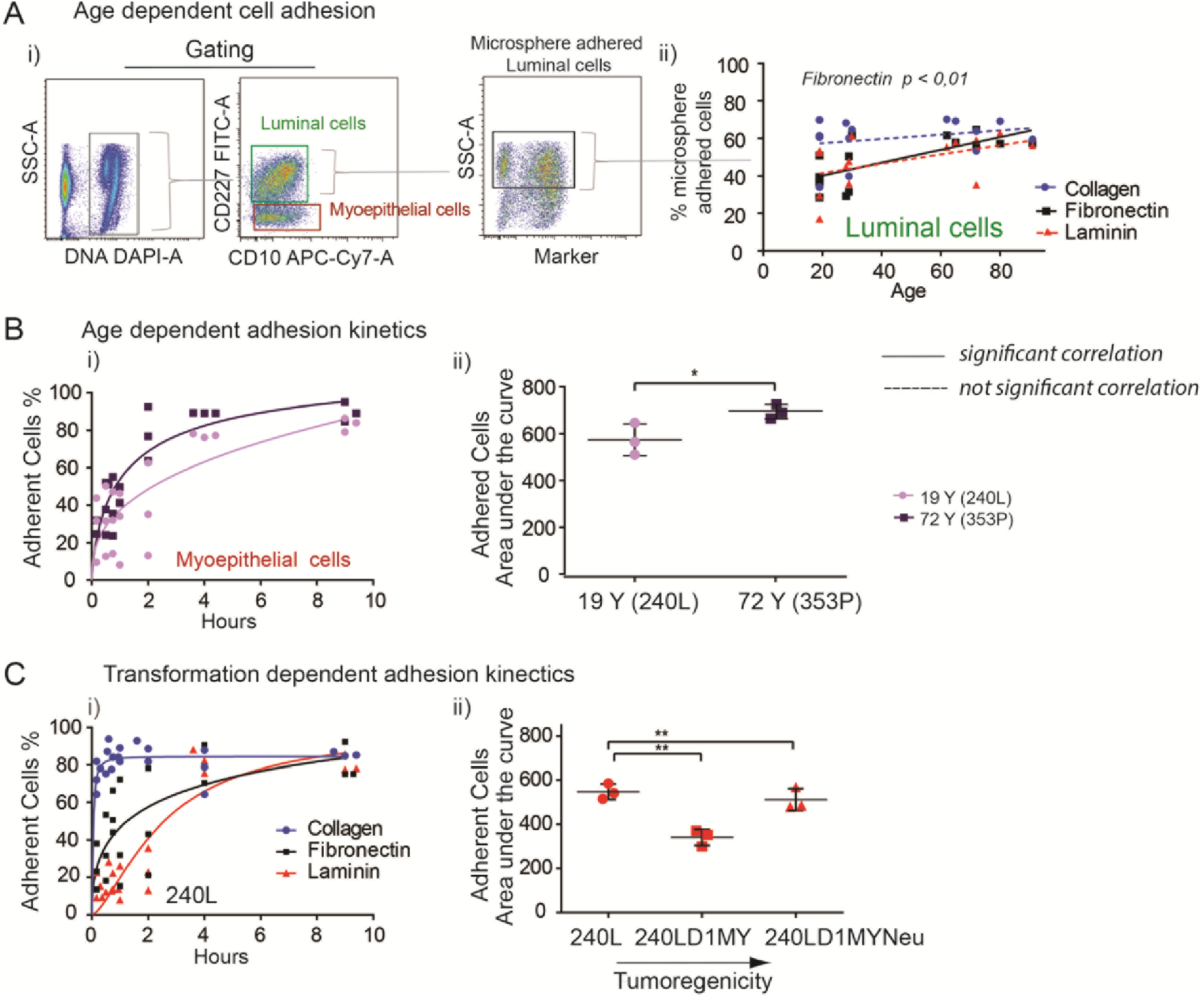
Cell adhesion efficiency of HMEC increase with age. (A) The percentage of cells adhered to different ECM-coated microspheres after 2.5 hours, as a function of age. HMEC from 10 women 19 – 91 y were included. i) Illustration of LEP-gating scheme (CD227^+^/CD10^-^). Side scatter values distinguishes cell-microsphere complexes from naked microspheres. ii) A significant linear regression correlation between age and cell adhesion abilities was found for LEP bound to FN. (B) Cell adhesion kinetics of HMEC from women 19 and 72 y (n=3 for each woman) to FN-coated microspheres was monitored for a period of 9 hours i) Hill’s curve of MEP+ cell adhesion.. ii) Comparison of AUC values of the Hill’s curves demonstrates significant difference in cell adhesion. (C) i) Cell adhesion kinetic analysis of HMEC (19 y, 240L) on different ECM. ii) Comparison of AUC-values of the cell adhesion kinetics to LAM-coated microspheres of an isogenic HMEC progression series (240L). The series comprised normal HMEC (240L), immortalized (240LD1MY) and oncogene-transformed (240LD1MYNeu) cells. The data demonstrate a significant decrease in cell adhesion with increasing tumorigenicity (n=3 for each cell type). Error bars are ±SD. P-values were calculated by Student’s two tailed t-test comparing AUC-values * P<0,05 ** = P<0,01.

**Suppl. Fig.5.**
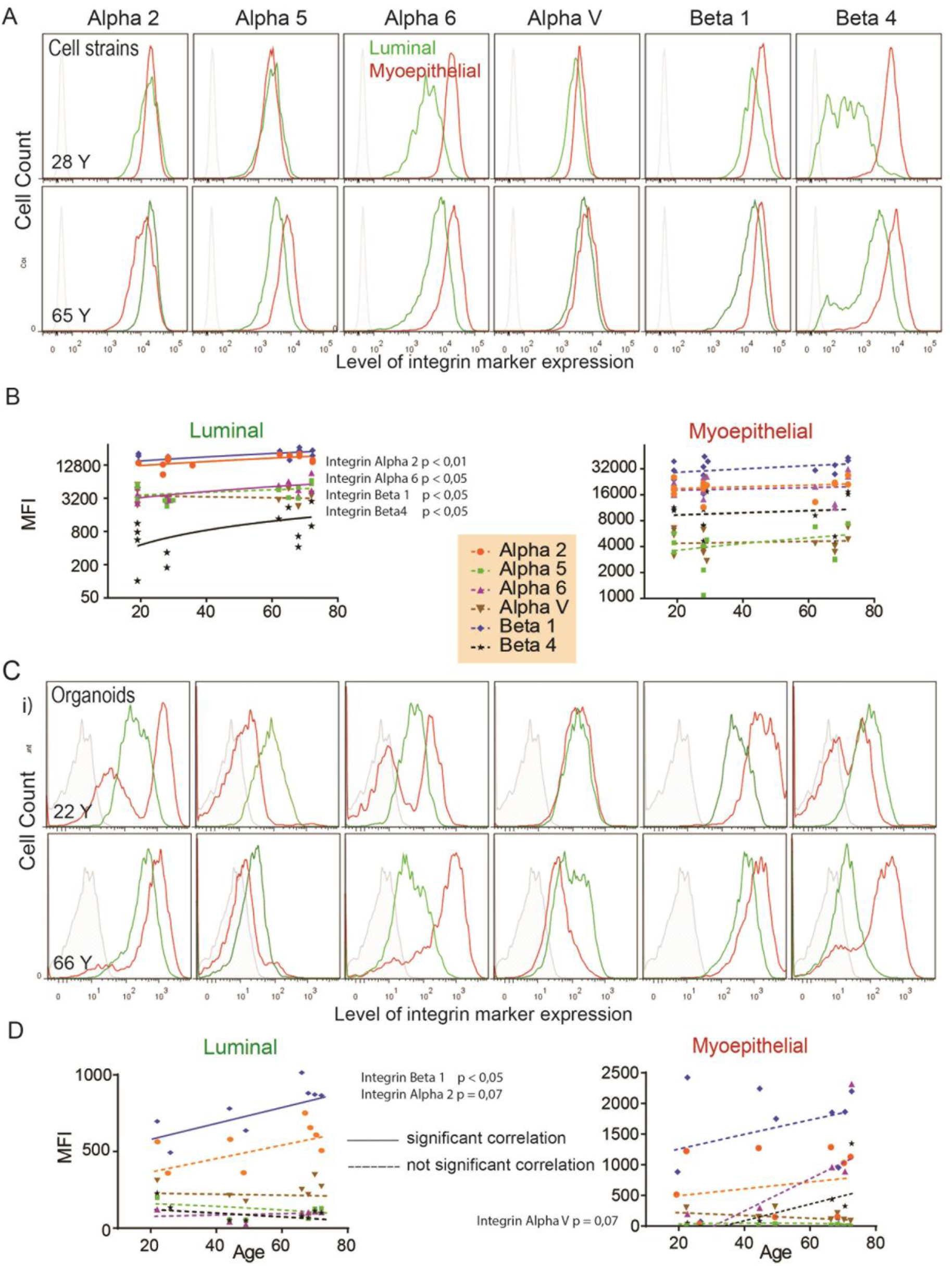
Integrin expression on HMEC increase with age. (A) Representative histograms from flow cytometry analysis of integrin expression in LEP and MEP from a 28 and a 65y woman. ITGα2, ITGα5, ITGα6, ITGFαV, ITGβ1 and ITGβ4 were measured. (B) Linear regression analysis of surface integrin levels on LEP and MEP from 7 women between 19 – 72 y (n=3 for each woman). (C) Representative histograms from flow cytometry analysis of integrin expression on LEP and MEP from non-passaged organoids of a 22 and 66y woman. (D) Linear regression analysis of surface integrin levels on LEP and MEP in non-passaged organoids from 9 women between 22 and 72y. Significant correlations (P<0.05) are demarcated in solid lines; non-significant correlations are represented by dotted lines.

**Suppl. Fig.6.**
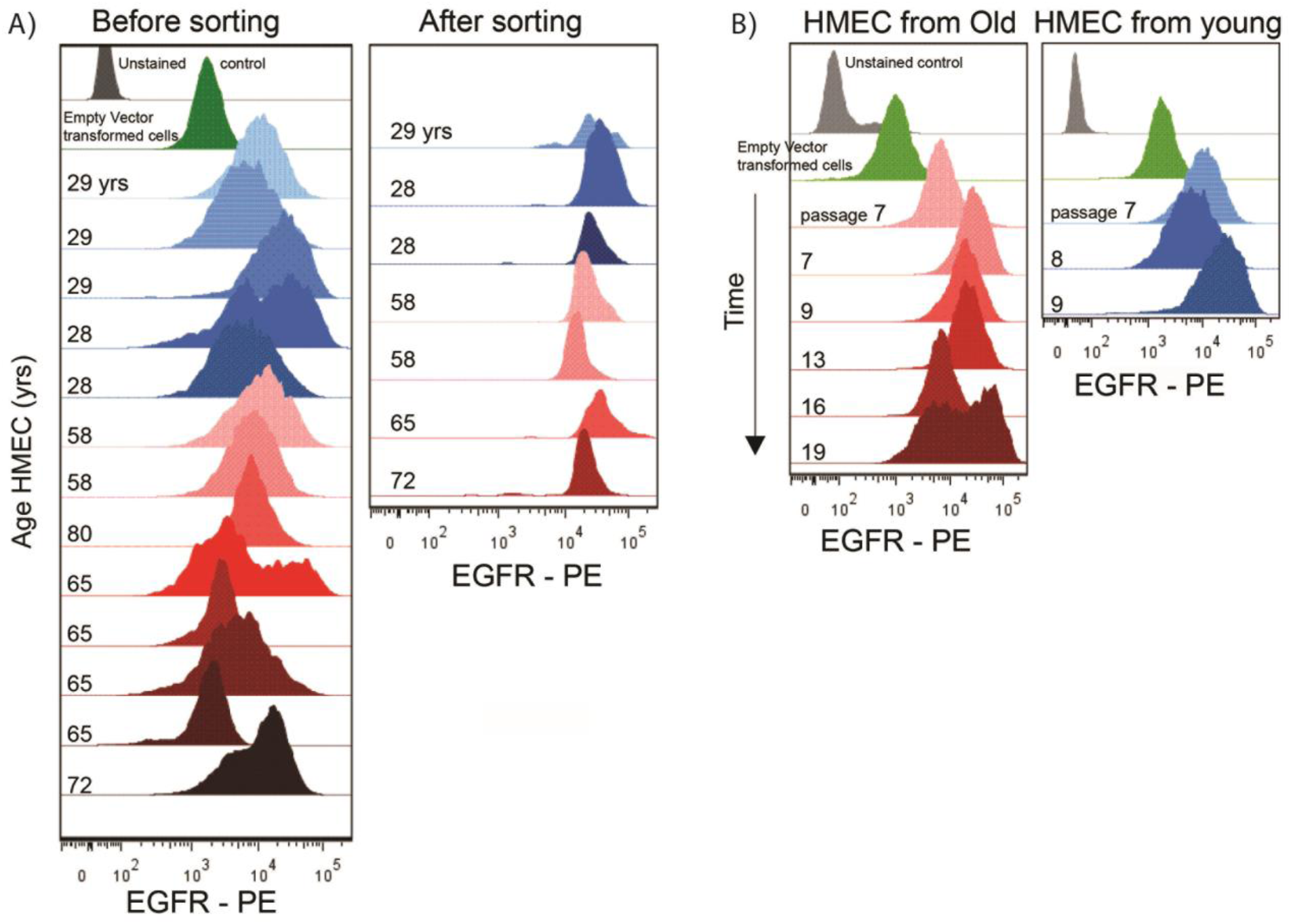
Post-sort EGFR^Del19^ oncogene expression is independent of HMEC age or passage. (A) FACS analysis of EGFR^Del19^ expressing HMEC. Post-sort HMEC EGFR^Del19^ expression was age-independent as HMEC from older women (red) and younger women (blue) showed similar levels. (B) FACS analysis of EGFR levels in EGFR^Del19^ expressing HMEC from an old (80 y) and young (29 y) woman during cell passage. EGFR expression levels appeared unchanged throughout passage of cells.

**Suppl. Fig.7.**
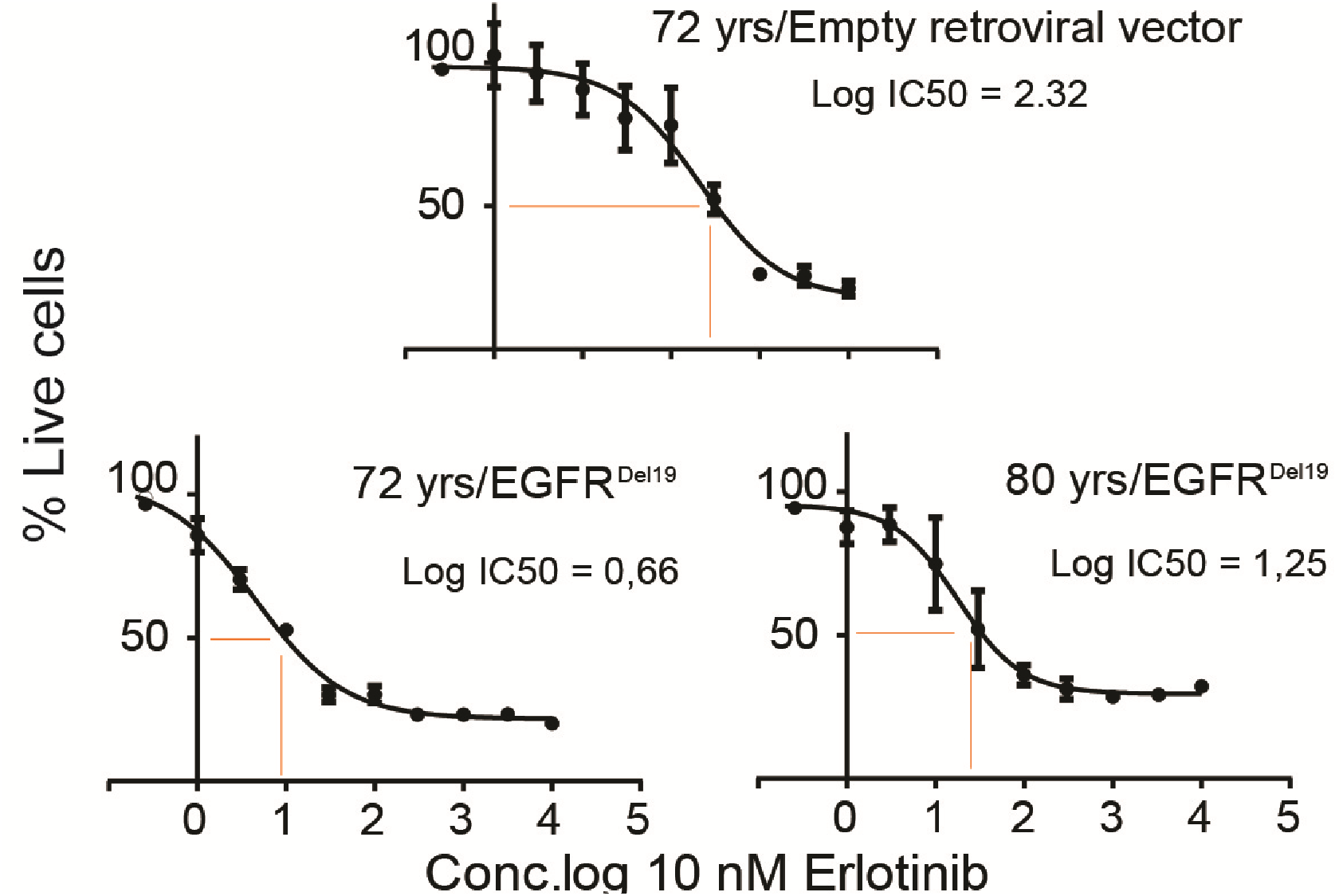
EGFR^Del19^ oncogene expressing cells are sensitive to Erlotinib. EGFR kinase inhibitor sensitivity of HMEC from two women (age 72 and 80 y) expressing EGFR^Del19^ or empty retroviral vector control. Non-linear regression analysis with variable slope shows the inhibitory effect of erlotinib on cell viability. Younger HMEC expressing EGFR^Del19^ became senescent precluding erlotinib sensitivity measure.

